# Efineptakin alfa (NT-I7) improves overall survival and induces tertiary lymphoid structures in murine lung tumors

**DOI:** 10.1101/2025.09.15.676444

**Authors:** Trang Dinh, Judong Lee, Sidra Islam, Namita Nanda, Dijana Bjelivuk, Dathan Andrews, Jiasen Zhang, Nikita L. Mani, Jin Zhou, Alexandra A. Wolfarth, Donghoon Choi, Rafi Ahmed, Joseph Skitzki, Deyu Fang, Weihong Guo, Zhenghe Wang, Rebecca C. Obeng

## Abstract

Tertiary lymphoid structures (TLSs) are emerging as good predictive biomarkers of response to cancer immunotherapy. However, therapeutic strategies to induce these structures are currently limited. We evaluated the therapeutic benefit of efineptakin alfa (NT-I7), a long-acting form of IL-7, and its ability to induce TLSs in a murine lung tumor model. NT-I7 improved overall survival in tumor-bearing mice. It also increased the abundance of T, B, dendritic cells, and stem-like CD8 T cells and promoted the formation of immune aggregates in the tumor microenvironment (TME). Stem-like CD8 T cells were preferentially located in the immune aggregates. Spatial transcriptomic analyses of the TME further demonstrated that the immune aggregates induced by NT-I7 included TLSs with enrichment of *Cd274* (PD-L1) transcripts and genes involved in antigen processing and presentation. Upregulation of *Cd274* in the TLSs may provide opportunities for synergy between NT-I7 and PD-1-targeted immunotherapy.

**STATEMENT OF SIGNIFICANCE:** This study demonstrates the ability of efineptakin alfa (NT-I7) to potentially augment the clinical efficacy of cancer immunotherapy by inducing tertiary lymphoid structures in the tumor microenvironment.

## INTRODUCTION

Tertiary lymphoid structures (TLSs) are ectopic lymphoid structures that develop in tissues and organs afflicted by chronic diseases such as cancer(1–3). These structures are composed of immune and stromal cells that morphologically and functionally resemble secondary lymphoid organs. A defining feature of TLSs is the tightly clustered T and B cells within the immune aggregates. The T and B cells can be organized into T- and B-cell-like zones as observed in the spleen and lymph nodes. Germinal centers can be observed in organized TLSs with evidence of polarization and somatic hypermutation(4, 5). The presence of TLSs within tumors has recently emerged as a critical determinant of response to immune checkpoint inhibitors (ICIs) such as blocking antibodies against the programmed cell death (PD-1) pathway(6–9). Therapeutic approaches to induce TLSs could improve clinical efficacy of cancer immunotherapy. However, the availability of such therapeutic agents is limited.

Interleukin-7 (IL-7) is a cytokine that supports naïve and memory T cell development, survival and proliferation, and the development of lymphoid organs(4, 10–12). IL-7 regulates lymphoid organogenesis by controlling the development and abundance of innate lymphoid cells that serve as lymphoid tissue inducer (LTi) cells(13, 14). LTi cells recruit and coordinate with lymphoid tissue organizer stromal cells for the development of lymphoid organs(15–17). Consequently, IL-7 and IL-7 receptor deficient mice lack LTi cells and have significant defects in the development of secondary lymphoid structures(18). In keeping with this, overexpression of IL-7 in mice leads to increased numbers of LTi cells, Payer’s patches, and functional TLSs(4). Thus, IL-7’s ability to support the development of TLSs could be therapeutically beneficial in combination with ICIs but its short half-life and inherent instability limits its use.

NT-I7 is a long-acting recombinant human IL-7 fusion protein (rhIL-7-hyFc) that is being investigated as a therapeutic option for patients with cancer(19). It has been shown to promote the expansion of T cells in murine models and in human studies(20, 21). In preclinical studies, NT-I7 has shown antitumor efficacy with increased intratumoral T cells(22–24) and supports the persistence of CAR T and CAR iNKT cells(21, 25, 26) in hematological and solid tumor models. However, its impact on TLS formation in cancer models has not been investigated. In this study, we determine the effect of NT-I7 on immune cell infiltration into tumors and characterize immune aggregates and TLSs induced by NT-I7.

## MATERIALS AND METHODS

### Mice and cell line

C57BL/6 mice were purchased from Jackson Laboratory. All experiments were performed in accordance with an approved Institutional Animal Care and Use Committee protocol. BalbC mice were purchased from Dae Han Bio Link (South Korea). All experiments were performed in accordance with National Institutes of Health guidelines, and protocols were approved by the Institutional Animal Care and Use Committee (IACUC) of POSTECH. The mice were randomized prior to treatment. The Lewis Lung Carcinoma cell line was a gift from the Ahmed and Kissick Labs at Emory University School of Medicine (Atlanta, GA). The cell line was authenticated prior to use and was routinely tested for contamination.

### Antibodies

CD45 (30-F11), CD3 (145-2C11), CD8a (53-6.7), CD4 (RM4-5), CD44 (IM7), PD-1 (29F.1A12), FOXP3 (MF-14), CD19 (6D5), B220 (RA3-6B2), MHC-II (M5/114.15.2), Gr-1 (RB6-8C5), CD11c (N418), CD11b (M1/70), and TIM-3 (RMT3-23) antibodies were purchased from BioLegend (San Diego, CA). TCF-1 (E6O1K) was purchased from Cell Signaling Technology (Danvers, MA) and Aqua live/dead was purchased from ThermoFisher (Oakwood, OH). For the CT26 model, primary antibodies were CD3 (145-2C11; Bio-Rad, Hercules, CA) and CD45R/B220 (RA3-6B2, Invitrogen, Waltham, MA) and secondary antibodies were goat anti-rat IgG (A11006) and goat ani-Armenian hamster IgG (A78966) purchased from ThermoFisher (Waltham, MA).

### Tumor growth and NT-I7 treatment

The lungs of seven-to ten-week-old male and female C57BL/6 mice were inoculated with 3 x 10^5^ LLC1 cells by tail vein injection. Treatment with 10 mg/kg of NT-I7 (NeoImmuneTech, Inc.) or control diluent by subcutaneous injection was initiated 7 days after tumor inoculation. The mice received one (n=19 per treatment group) or two doses (n=8 per treatment group) 7 days apart. Tumor-bearing lungs were harvested between days 16 and 25 after tumor inoculation or at the time of death for analysis. The dispositions of the mice were recorded and used for the survival analysis.

For the colorectal model, seven-week-old female BalbC mice were intraperitoneally (i.p.) implanted with 3 x 10^5^ CT26 cells to initiate disseminated tumor at day 0. The mice received a single intramuscular injection of NT-I7 (10 mg/kg) on day 7. CT26 tumors were harvested 10 days after the treatment, fixed in formalin and paraffin-embedded prior to immunohistochemistry staining.

### Flow cytometry

Tumor infiltrating immune cells were enriched by percoll (Cytiva, Europe) gradient separation after tumor digest with collagenase IV (Worthington Biochemical, Lakewood, NJ) and then used for flow cytometric analysis. Briefly, the single cell suspensions were incubated with surface antibodies at room temperature, then fixed and permeabilized with the eBioscience Foxp3/Transcription factor staining buffet set (ThermoFisher), and then finally incubated with intracellular antibodies. The samples were processed on the BDFortessa instrument and analyzed using FlowJo (Treestar, v10).

### Tissue histology and Immunofluorescence

Sections of tumor samples were formalin-fixed and embedded in paraffin for histologic analysis. 5-micron sections were cut and stained with hematoxylin and eosin. Additional tumor samples were frozen in optimal cutting temperature (OCT) medium and 8-micron sections of the tumor were stained with antibody markers of interest for immunofluorescence. The slides were imaged using a Leica HyVolution SP8 confocal microscope and the images were analyzed using QuPath (v. 0.5) and ImageJ.

### Immunohistochemistry

5-micron sections of formalin fixed paraffin embedded peritoneal CT26 tumors were deparaffinized, rehydrated, and subjected to antigen retrieval prior to incubating with primary antibodies (CD45R/B220 and CD3). Following incubation with primary antibodies, the sections were incubated with fluorescently tagged secondary antibodies. Images of the stained sections were acquired on confocal and analyzed using Zeiss Software (ZEN 3.3 Lite).

### Spatial transcriptomics

The filtered count matrices and spatial metadata generated by Space Ranger were analyzed using the Seurat R package (v5.2.0) within R (v4.4.1). Data normalization was performed using the SCTransform() and RunPCA() was conducted for linear dimension reduction. The top 30 principal components were further utilized for non-linear dimension reduction via RunUMAP and for FindNeighbors(). Unsupervised cell type clustering was performed using the Louvain algorithm via FindClusters() functions with a resolution parameter of 1.2. Top gene markers for all the clusters were obtained using the Wilcoxon rank-sum test implemented in FindAllMarkers() with min.pct of 0.1 and logfc.threshold = 0.25.

Supervised cell type clustering based on single-cell reference mapping was performed using combined scRNAseq references from the Tabula Muris Consortium and from Kumar et al via FindTransferAnchors() and TransferData() functions with CCA reduction. Prediction scores across tissue maps for each cell type cluster were visualized via SpatialFeaturePlot(). Differential expression analysis between the NT-I7 treated group versus the control group was performed via FindMarkers(). Then gene set enrichment analyses (GSEA) were conducted on the ranked differential gene list using clusterProfiler’s GSEA() with the mouse hallmark geneset from mSigDB and Benjamini-Hochberg correction for p-values. All spatial plots and statistical graphics were generated using Seurat’s SpatialDimPlot, SpatialFeaturePlot, DimPlot, FeaturePlot, ggplot2, cowplot, pheatmap, and patchwork.

### Statistical analysis

Continuous variables were first assessed for normal distribution using four different tests (D’Agostino-Pearson omnibus normality, Anderson-Darling, Shapiro-Wilk, and Kolmogorov-Smirnov with Dallal-Wilkinson-Lilliefor P value). Unpaired t test was used to evaluate differences between groups for data with a normal distribution. Mann-Whitney test was used to evaluate the differences between groups for data that did not have a normal distribution. Paired t test was performed similarly when comparing groups within the same samples. The Binomial test was used to compare observed and expected proportions. Statistical analyses were performed using GraphPad Prism 10 (v10.4.1) and R.

## RESULTS

NT-I7 is associated with improved survival and increased numbers of immune cells in the tumor microenvironment IL-7 signaling promotes proliferation and survival of T cells(10). To determine the impact of NT-I7 on tumor control and tumor infiltrating immune cells, we seeded the lungs of mice with LLC1 cells. Seven days after tumor inoculation, the mice were treated once a week with a total of one (n=5-6 per group) or two doses (n=14 per group) of NT-I7. The mice were monitored over time for survival. A significant improvement in overall survival was observed in tumor-bearing mice treated with NT-I7 (Fig. 1A). Tumors were also harvested between days 16 and 25 after tumor inoculation to evaluate the abundance of immune cells in the tumors by flow cytometry. The gating scheme is shown in figure 1B. The number of CD45^+^ cells per gram of tumor tissue significantly increased after NT-I7 treatment (Fig. 1C). Significantly more B cells and CD8 T cells were also observed in the NT-I7-treated group (Fig. 1D,E). No differences in the number of antigen-experienced CD8 T cells per gram of tumor tissue was observed between the two groups (Fig. 1F). We also observed an increase in the number CD4 T cells in the NT-I7 treated tumors (Fig. 1G). However, there was no difference in the number of regulatory CD4 T cells per gram of tumor tissue between control and NT-I7 treated mice (Fig. 1H). While no difference in CD11b^+^CD11c^+^ myeloid cells was noted, a significant increase in conventional dendritic cells (cDCs) was evident (Fig. 1I). Further analysis of the CD44^+^PD-1^+^ antigen experienced CD8 T cell population demonstrated a significant decrease in the proportion of terminally differentiated TIM-3^+^ cells (Fig. 2A). Interestingly, the proportion of stem-like CD8 T cells, marked by positive TCF-1 and negative TIM-3 expression, within the CD44^+^PD-1^+^ population was significantly increased after NT-I7 therapy.

**Figure 1.**
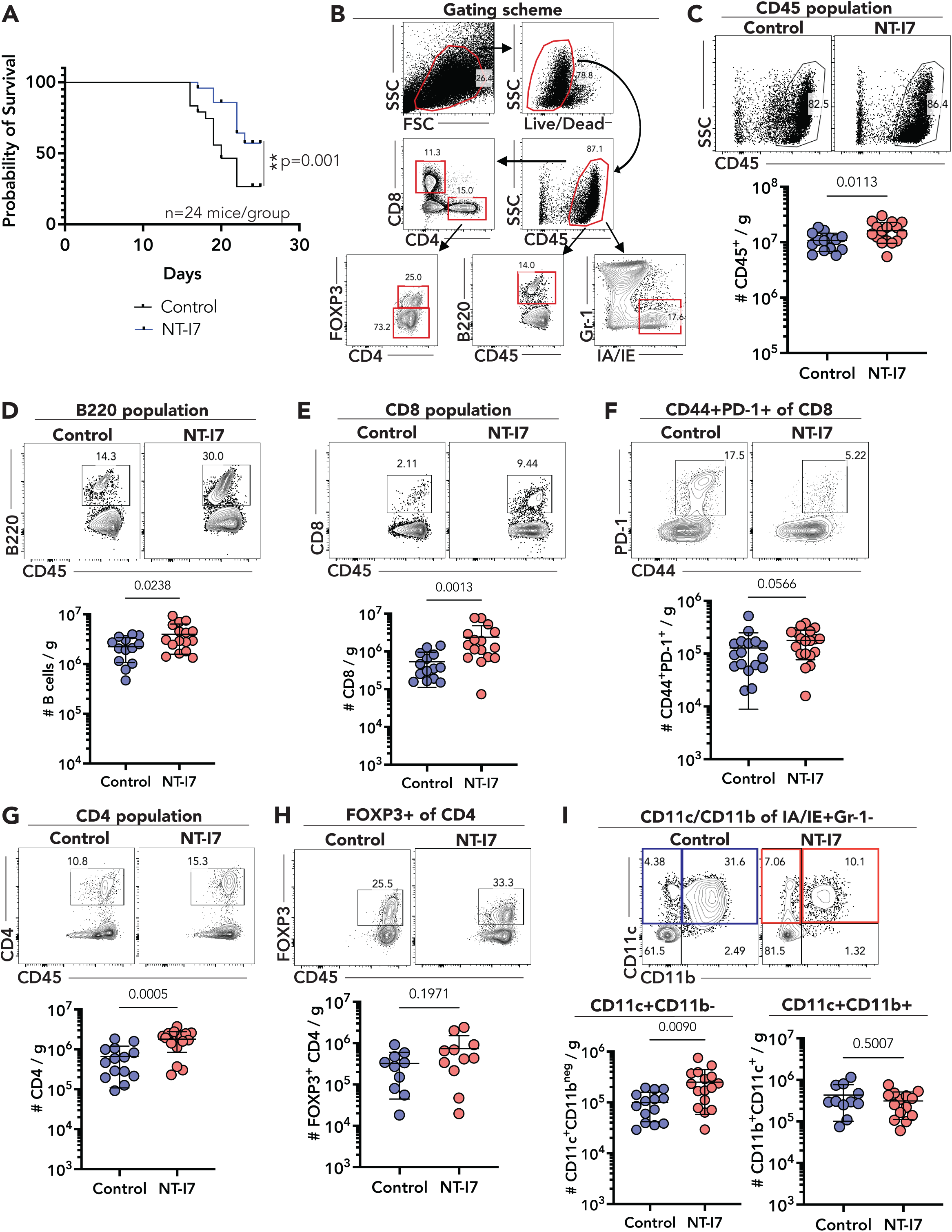
Overall survival and tumor immune cell infiltrates in NT-I7-treated mice. (A) Kaplan-Maier survival curve for control and NT-I7-treated mice (n=24 per group). (B) Representative dot plots of gating scheme. (C-I) Representative dot plot and summary graphs for CD45+ (C), B cells (D), CD8 T cells (E), activated CD8 T cells (F), CD4 T cells (G), T regulatory cells (H), and CD11c and CD11b cell subsets (I). N=10-19 (control) and 11-20 (NT-I7) mice per group. Unpaired t test (A, B, F, and H (CD11c+CD11b-)) and Mann-Whitney test (C, D, G, and H (CD11c+CD11b+)) were performed for data with normal distribution and non-normal distribution respectively. Horizontal lines represent the mean and error bars represent standard deviation. FSC – forward scatter; SSC – side scatter.

**Figure 2.**
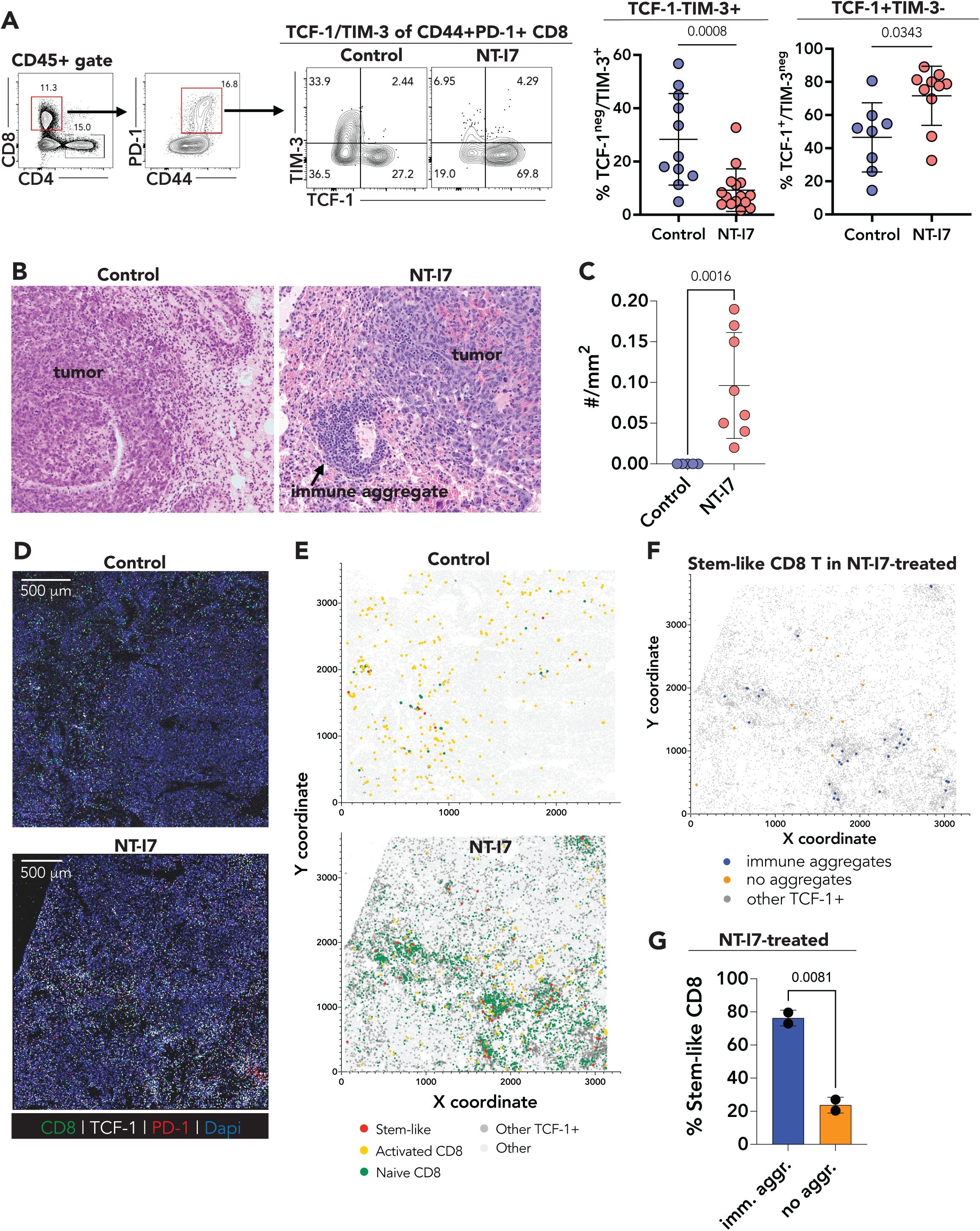
Stem-like CD8 T cell abundance and localization in NT-I7-treated tumor tissues. (A) Representative dot plot and summary graphs for effector CD8 (TCF-1-TIM-3+; n=11 – 14 mice per group) and stem-like CD8 (TCF-1+TIM-3-; n=8 – 10 mice per group) subsets of activated CD8 T cells. (B) Representative H&E images of LLC-1 tumors in the lungs. (C) Summary plot of manual counts of immune aggregates in tumor-bearing lung tissues per mm^2^. Each datapoint represents average #/mm^2^ per sample (n=5 for control and 8 for NT-I7-treated samples). (D) Representative images of immunofluorescently stained tumor-bearing lungs from control and NT-I7-treated mice. (E) Representative 2D representation of immune cells in control and NT-I7 treated tumors (n=2 per group). (F and G) Representative plot (F) and summary graph (G) of location of stem-like CD8 T cells in NT-I7-treated tumors from E (n=2). Mann-Whitney test (A and C) and unpaired T test (G). Horizontal lines represent the mean and error bars represent standard deviation. Imm. – immune; aggr. – aggregates.

### Stem-like CD8 T cells are localized in immune aggregates induced by NT-I7

We have previously demonstrated that stem-like CD8 T cells are preferentially located in tertiary lymphoid structures that are present in the tumor microenvironment and metastatic lymph nodes of human tumors(27, 28), and in secondary lymphoid organs in the murine lymphocytic choriomeningitis virus (LCMV) chronic viral infection model(29). We therefore examined the tissues of control and NT-I7-treated mice for the presence of immune aggregates and sought to determine the location of the stem-like CD8 T cells in the tumor microenvironment. On H&E sections of control mice, no lymphoid aggregates were observed (Fig. 2B,C). In contrast, immune aggregates abutting tumor nodules and within the normal lung parenchyma were observed in NT-I7 treated mice. An average of 0.096 immune aggregates per mm^2^ were identified in NT-I7-treated tumor-bearing tissues (Fig. 2C).

Immunofluorescence was performed on the tissue sections to identify subsets of CD8 T cells including stem-like CD8 T cells. NT-I7-treated tissues had increased infiltration of TCF-1^+^ lymphoid cells including naïve and naïve-like (CD8^+^ TCF-1^+^ PD-1^neg^) and stem-like (CD8^+^ TCF-1^+^ PD-1^+^) CD8 T cells (Fig. 2D,E). Consistent with our flow analysis, more stem-like CD8 T cells were present in the NT-I7-treated tumors compared to the control tissue samples by immunofluorescence. Using TCF-1^+^ cell clusters as a surrogate marker for immune aggregates, we mapped the location of stem-like CD8 T cells (Fig. 2E) and assessed their proximity to the TCF-1^+^ immune aggregates (Fig. 2F). TCF-1^+^ aggregates of varying sizes were observed in the NT-I7-treated tissues with only rare aggregates seen in the control tissues. Seventy-three to 79% of the stem-like CD8 T cells were localized within or in close proximity of the immune aggregates (Fig. 2G and Table 1) indicating that the stem-like CD8 T cells may preferentially home to immune aggregates within the tumor microenvironment.

**Table 1.** Stem-like CD8 T cells are preferentially located in TLSs

| Replicate 1 |  |  |  |  | Replicate 2 |  |  |  |
| --- | --- | --- | --- | --- | --- | --- | --- | --- |
|  | Expected (%) | Observed (%) | 95% CI of Observed | p-value | Expected (%) | Observed (%) | 95% CI of Observed | p-value |
| TLS | 24 (50) | 35 (72.92) | 59 – 83.43 | **0.002**<br>(2-tailed) | 56.5 (50) | 90 (79.65) | 71.31 – 86.04 | **<0.0001**<br>(2-tailed) |
| Non-TLS | 24 (50) | 13 (27.08) | 16.57 – 41 |  | 56.5 (50) | 23 (20.35) | 13.96 – 28.69 |  |

Binomial test comparing observed vs. expected outcomes using Wilson/Brown method to calculate confidence intervals (CI)

### Immune aggregates induced by NT-I7 include TLSs ranging from unorganized to well-organized TLSs

Upon further characterization of the tumor sections by immunofluorescence, we observed that T and B cells were mainly scattered throughout the tumors of the control mice (Fig. 3A). Majority of the tumors from control mice lacked aggregates of lymphocytes. A few small clusters of T cells only and a couple of small clusters containing both T and B cells were observed in three out of the eight control tumors (Fig. 3B,C). B cell-rich clusters were not identified in the control tissues. In contrast, we observed significantly more lymphocyte aggregates in the NT-I7 treated tumors (Fig. 3A,B). These aggregates consisted of T cell-rich, B cell-rich, or a mixture of T and B cells indicative of TLSs (Fig. 3D). On average, B cell-rich aggregates accounted for 5% of the immune aggregates while T cell-rich aggregates and aggregates containing both T and B cells made up 24% and 71% of the aggregates in the NT-I7-treated tissues respectively. The immune aggregates were also larger in the NT-I7-treated tumors (Fig. 3E). Some of the T-and-B cell aggregates in the NT-I7-treated tumors were organized into distinct T-and B-cell-like zones, similar to the organization in secondary lymphoid organs suggesting potential functionality of the TLSs (Fig. 3D). The TLSs (T-and-B cell aggregates) in the NT-I7-treated tumors were also significantly larger than the T cell-rich and B cell-rich aggregates (Fig. 3F). We observed similar findings in a CT26 colorectal tumor model (Suppl. Fig. 1).

**Figure 3.**
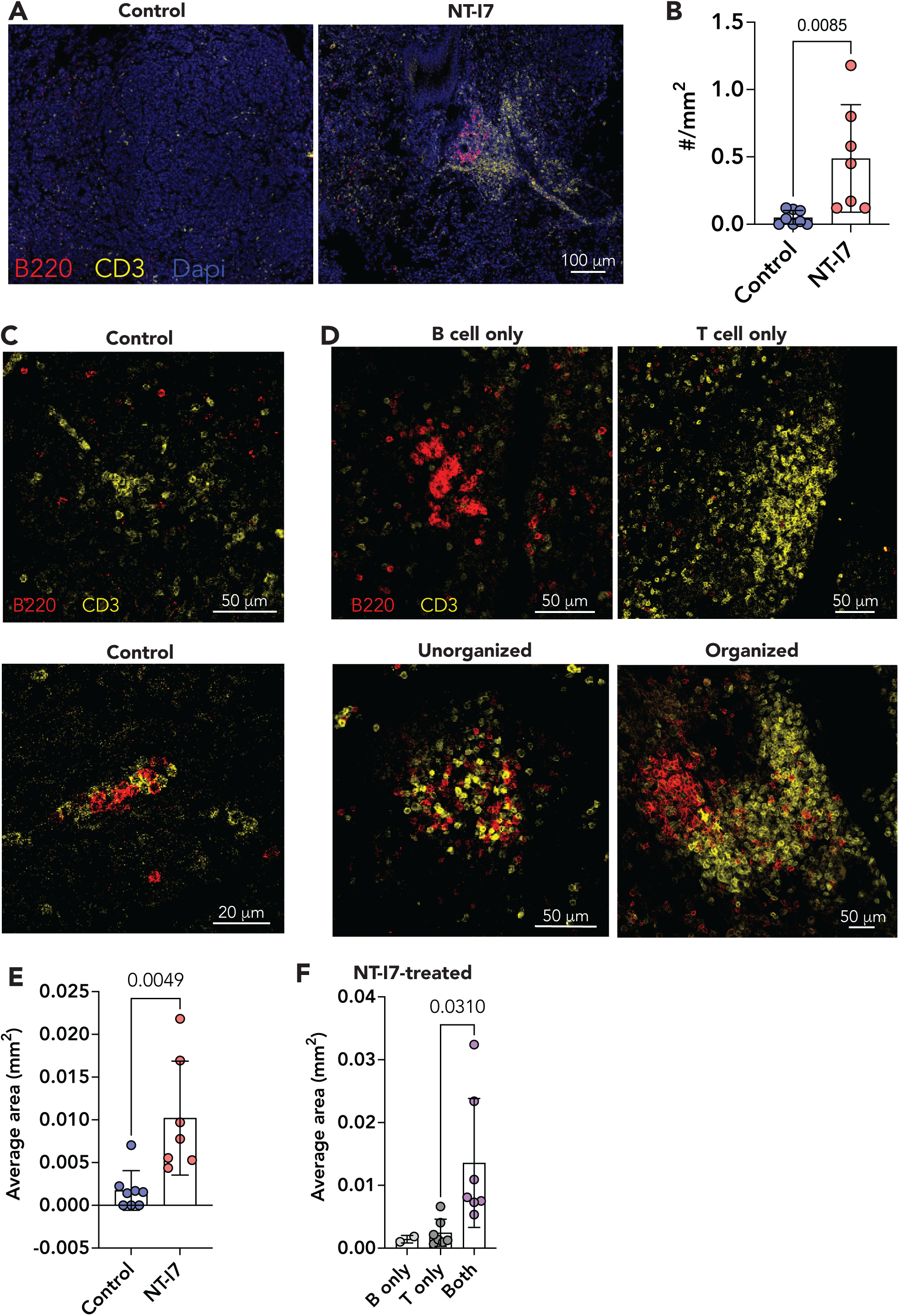
Presence and types of immune aggregates in control and NT-I7-treated tumor samples. (A) Representative IF images of control and NT-I7-treated samples. (B) Summary plot of manual counts of immune aggregates in tumor-bearing lung tissues per mm^2^. Each datapoint represents average #/mm^2^ per sample (n=8 for control and 7 for NT-I7-treated samples). (C and D) Representative IF images of immune aggregates identified in control and NT-I7-treated samples. (E and F). Summary plot of the average area of aggregates per sample (E) or aggregate type (F); n=8 for control and 7 for NT-I7-treated samples.

### Spatial transcriptomic analysis of immune cell infiltrates and TLSs after NT-I7 treatment

We performed spatial transcriptomic analysis using the 10x Visium platform to further examine the composition of the immune aggregates induced by NT-I7. We first distinguished the tumor regions from the normal lung parenchyma (Fig. 4A-B). Unsupervised clustering identified 16 unique clusters (Fig. 4C and Suppl. Table 1). UMAP visualization suggested an underrepresentation of cluster 13, and an overrepresentation of clusters 2 and 6 in the NT-I7-treated tissue (Fig. 4D, Table 2, and Suppl. Table 1). Using the FindAllMarkers function of Seurat to further analyze the clusters, we determined that clusters 2 (epithelial/immune mix), 6 (lymphocytes/dendritic cells), 7 (macrophages/myeloid cells), and 13 (alveolar macrophages) were enriched for genes of hematopoietic lineage (Fig. 4E-F). Differential gene set expression analysis revealed enrichment of cell lineage gene transcripts such as *Cd3d*, *Ptprc*, *Cd79a*, *Lamp3*, and *Cd163* suggesting an enriched heterogenous immune population of T, B, and myeloid cells in the NT-I7-treated tumors (Fig. 4G,H). Genes involved in antigen processing and presentation such as *Cd74*, *Cd84*, *H2-Ab1*, and *H2-Aa* were also upregulated in the NT-I7-treated tumor. Notably, coinhibitory transcripts like *Cd274* (PD-L1), *Btla*, and *Cd200* were not upregulated in the NT-I7-treated tumor. Gene transcripts that define adhesion molecules such as *Sell*, which directs lymphocyte migration, as well as *Icam2* that regulates T cell adhesion and migration were enriched in the NT-I7-treated tumor. Additionally, chemokines that support migration of immune cells like *Cxcl3*, *Ccl5*, and *Spp1* were enriched. The *Ccr7* chemokine receptor that directs migration of T cells into lymphoid tissues was also upregulated. The data suggest that IL-7 signaling promotes migration and organization of immune cells in the tumor microenvironment.

**Figure 4.**
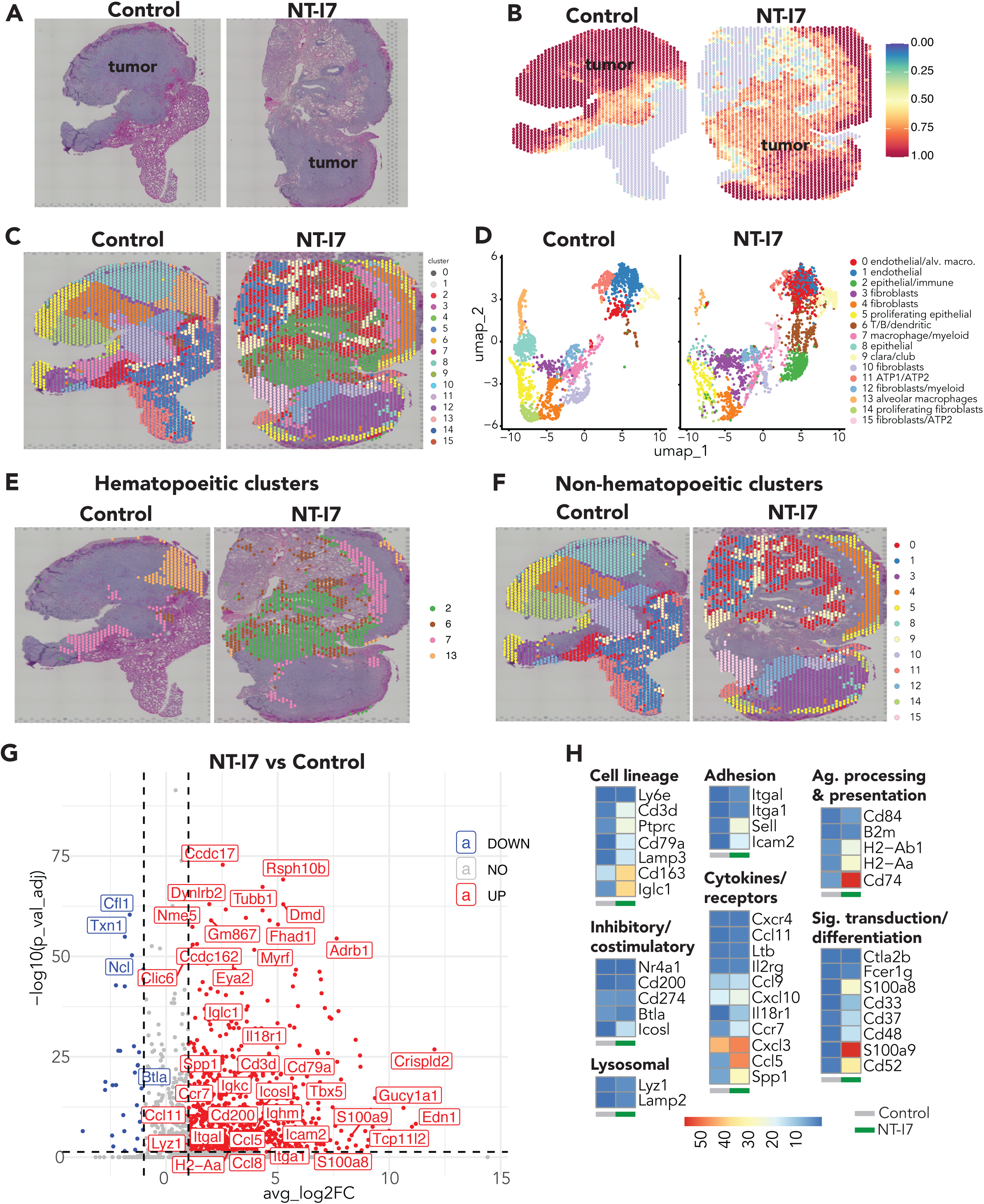
Spatial transcriptomic analyses of tumor samples from control and NT-I7-treated mice. (A and B) H&E images (A) and spatial representation of tumor regions (B) in control and NT-I7 treated tumor samples. (C, E, F) Spatial localization of unsupervised clusters. (D) UMAP of unsupervised clusters from control and NT-I7-treated samples. (G) Volcano plot of differentially expressed genes. (H) Heatmap of selected immune-related gene transcripts. Alv. macro. – alveolar macrophages; ATP – alveolar type pneumocytes.

**Table 2.**
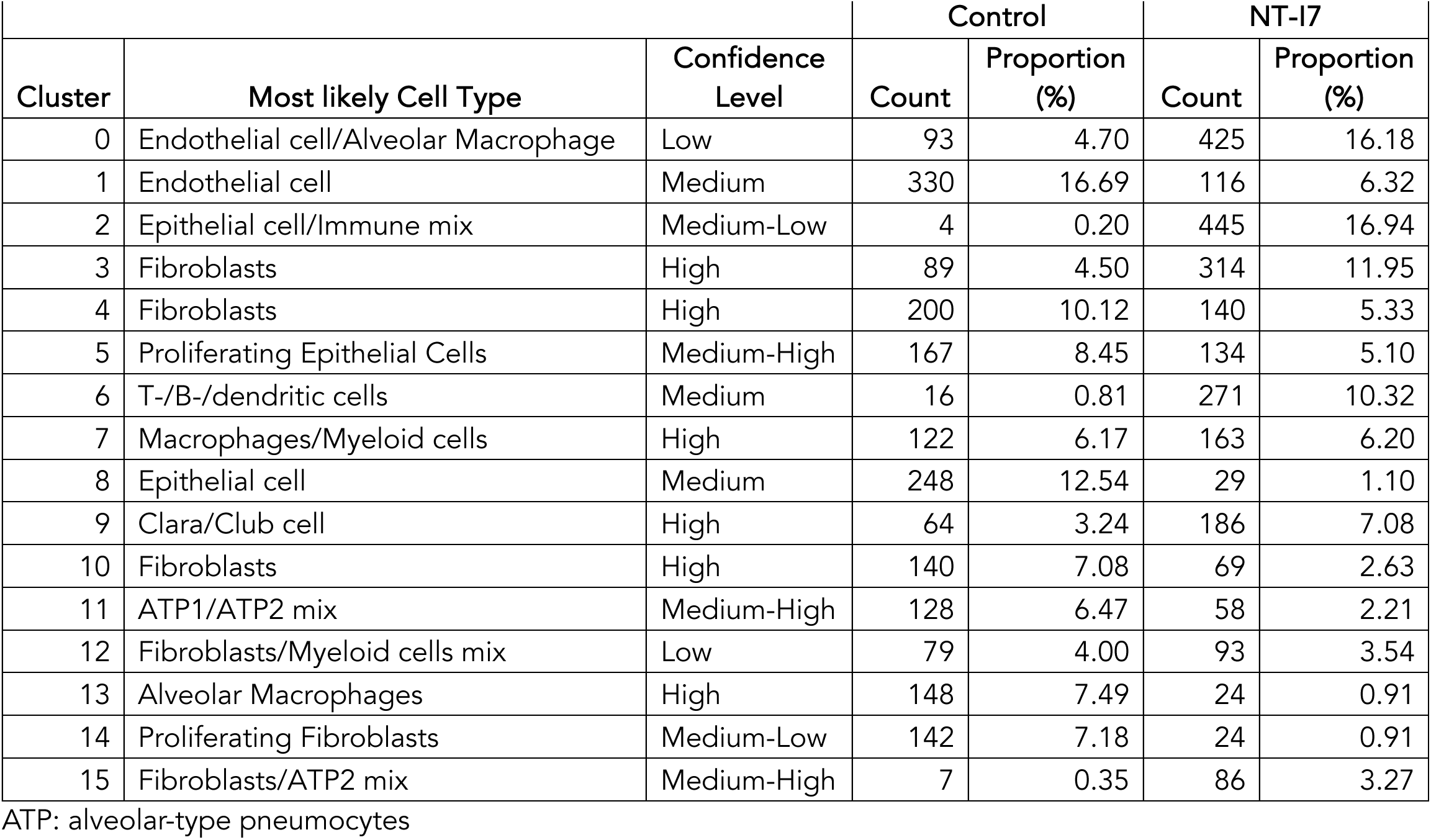
Unsupervised clusters with annotated cell types and abundance in tumors

|  |  |  | Control |  | NT-I7 |  |
| --- | --- | --- | --- | --- | --- | --- |
| Cluster | Most likely Cell Type | Confidence Level | Count | Proportion (%) | Count | Proportion (%) |
| 0 | Endothelial cell/Alveolar Macrophage | Low | 93 | 4.70 | 425 | 16.18 |
| 1 | Endothelial cell | Medium | 330 | 16.69 | 116 | 6.32 |
| 2 | Epithelial cell/Immune mix | Medium-Low | 4 | 0.20 | 445 | 16.94 |
| 3 | Fibroblasts | High | 89 | 4.50 | 314 | 11.95 |
| 4 | Fibroblasts | High | 200 | 10.12 | 140 | 5.33 |
| 5 | Proliferating Epithelial Cells | Medium-High | 167 | 8.45 | 134 | 5.10 |
| 6 | T-/B-/dendritic cells | Medium | 16 | 0.81 | 271 | 10.32 |
| 7 | Macrophages/Myeloid cells | High | 122 | 6.17 | 163 | 6.20 |
| 8 | Epithelial cell | Medium | 248 | 12.54 | 29 | 1.10 |
| 9 | Clara/Club cell | High | 64 | 3.24 | 186 | 7.08 |
| 10 | Fibroblasts | High | 140 | 7.08 | 69 | 2.63 |
| 11 | ATP1/ATP2 mix | Medium-High | 128 | 6.47 | 58 | 2.21 |
| 12 | Fibroblasts/Myeloid cells mix | Low | 79 | 4.00 | 93 | 3.54 |
| 13 | Alveolar Macrophages | High | 148 | 7.49 | 24 | 0.91 |
| 14 | Proliferating Fibroblasts | Medium-Low | 142 | 7.18 | 24 | 0.91 |
| 15 | Fibroblasts/ATP2 mix | Medium-High | 7 | 0.35 | 86 | 3.27 |

ATP: alveolar-type pneumocytes

We then performed supervised clustering to identify specific cell lineages based on single-cell reference mapping with combined scRNAseq references from the Tabula Muris Consortium (Lung datasets) and from Kumar et al(30) (Fig. 5A,B). Consistent with our flow cytometry analysis (Fig. 1) and the analysis of the unsupervised clusters (Fig. 4), T and B cell-related gene transcripts within the immune regions were more abundant in the NT-I7-treated tissue while the myeloid-related transcripts were increased in the control tumor (Table 3). As observed with the unsupervised clustering, gene transcripts of several cell lineages (*Cd3d*, *Ptprc*, Lamp3, and *Cd163*) were upregulated in the NT-I7-treated tumor (Fig. 5C). Mapping the spatial location of different immune subsets showed that the T, B, and myeloid cells were located mainly at the periphery of the tumor bed in the control and NT-I7 treated tumor (Fig. 5D). However, focal areas with increased levels of T and B cell-related genes were present in the NT-I7-treated tumor. These areas mapped to immune aggregates identified in the corresponding H&E images of the NT-I7-treated tumor (Fig. 5D).

**Figure 5.**
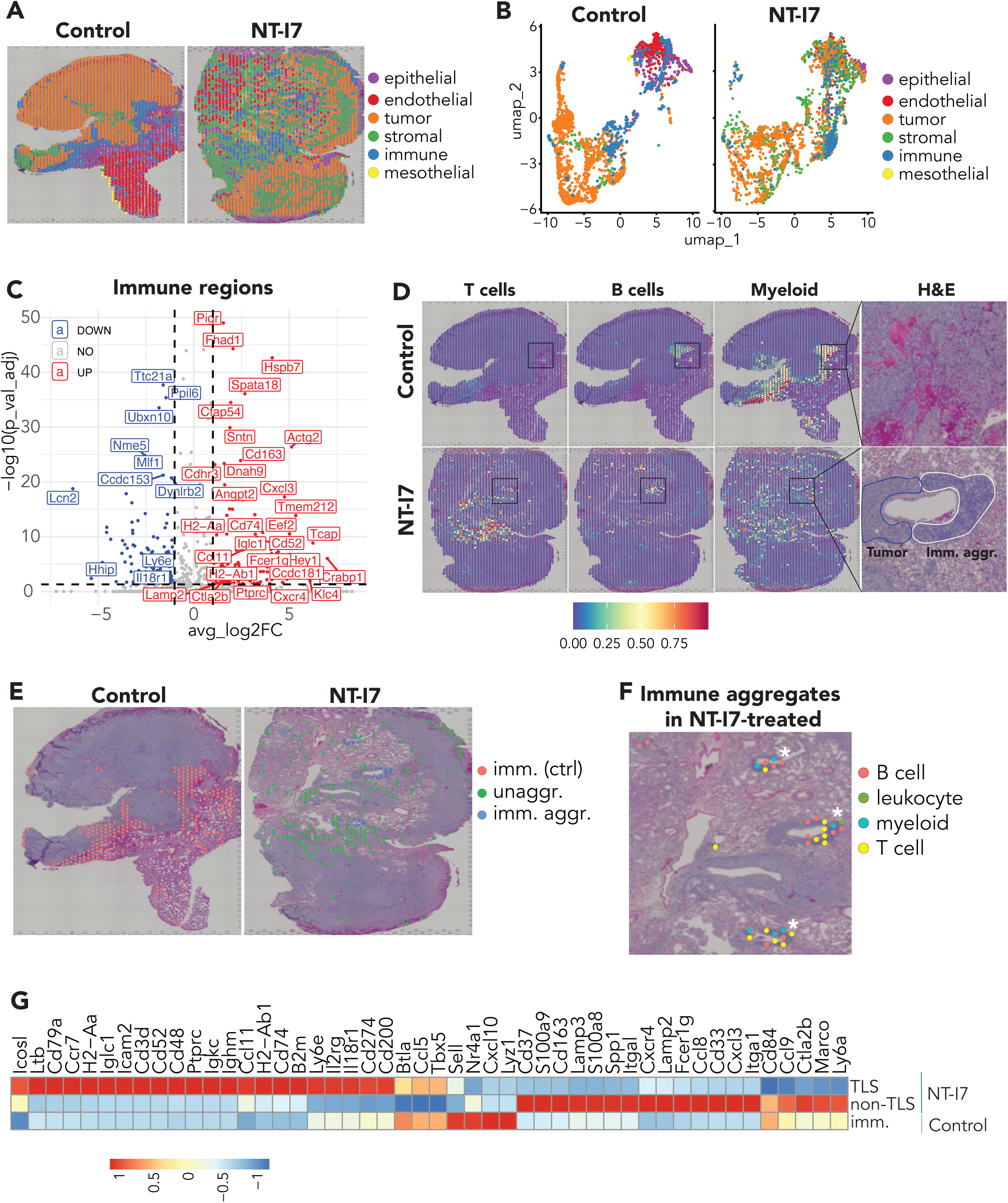
Transcriptional profile of immune aggregates including TLSs induced by NT-I7. (A,B) Spatial representation and UMAP of supervised clusters. (C) Volcano plot of differentially expressed genes from the immune population. (D) Spatial localization of cell lineage specific gene transcripts for T, B, and myeloid cells in relation to H&E profile. (E) Spatial localization of aggregated and unaggregated immune regions. (F) Spatial localization and lineage identities of immune aggregate regions in the NT-I7-treated sample. White asterisks indicate aggregates that are TLSs. (G) Heatmap of the transcriptional profile of TLS and non-TLS regions in the NT-I7-treated tumor and the immune regions in the control tumor. Ctrl – control; unaggr. – unaggregated.

**Table 3.** Counts and proportion of supervised clusters

| Cluster | Control |  | NT-I7 |  |
| --- | --- | --- | --- | --- |
|  | Count | Proportion (%) | Count | Proportion (%) |
| T/B cell | 2 | 0.1 | 131 | 4.99 |
| endothelial cell | 305 | 15.43 | 256 | 9.74 |
| epithelial cell | 167 | 8.45 | 105 | 4.00 |
| leukocyte | 128 | 6.47 | 93 | 3.54 |
| LLC Tumor | 1098 | 55.54 | 871 | 33.16 |
| macrophage | 147 | 7.44 | 83 | 3.16 |
| mesothelial cell | 18 | 0.91 | 1 | 0.04 |
| stromal cell | 112 | 5.67 | 1087 | 41.38 |

We focused specifically on the immune regions from the supervised clustering to identify the location of the immune cells in control and NT-I7-treated tumors and separated the immune regions into immune aggregate and unaggregated regions (Fig. 5E). T cell-specific lineage genes were enriched in some of the immune aggregates while others were enriched for B cell lineage genes (Fig. 5F). Other aggregates exhibited enrichment of genes corresponding to multiple cell lineages including B cell, T cell, and myeloid-specific lineages, consistent with TLSs. Comparison of the gene transcripts from the immune regions between the control tissues and the TLS and non-TLS regions of the NT-I7 revealed enrichment of negative immune regulators (*Btla* and *Nr4a1*) and some chemokines and adhesion genes (*Cxcl10* and *Sell*) in the immune regions from the control tissue (Fig. 5G). Enriched genes in the non-TLS immune regions in the NT-I7-treated tissues included chemokines and receptors (*Ccl9*, *Ccl8*, *Cxcl3*, and *Cxcr4*), transcripts involved in signal transduction and differentiation (*Cd33*, *Cd37*, *S100a8*, *S100a9*, *Ctla2b*, and *Fcer1g*), and cell lineages such as histiocytes (Cd163) and dendritic cells (Lamp3). Within the TLSs in the NT-I7-treated tissues, *Ltb*, a cytokine known to drive lymphoid neogenesis(31, 32), and other genes that regulate immune cell migration into lymphoid tissues (*Ccl11*, *Ccr7*), adhesion (*Icam2*), and those involved in antigen processing and presentation (*B2m*, *Cd74*, *H2-Aa*, *H2-Ab1*) were enriched. We also observed enrichment of the costimulatory gene *Icosl*. In contrast to our comparison of the whole TME between control and NT-I7 treated tumors (Fig. 4), coinhibitory gene transcripts such as *Cd274* (PD-L1) and *Cd200* were enriched within the TLSs induced by NT-I7 but not in the non-TLS regions and the immune regions in the control tumor. Interestingly, *Igkc*, an immunoglobulin that has been associated with progression-free survival in solid tumors(33) was among the top differentially expressed genes in the TLSs induced by NT-I7.

## DISCUSSION

The presence of TLSs in the tumor microenvironment is emerging as a predictive biomarker for durable response to cancer immunotherapy(6–8, 34). However, therapeutic strategies aimed at inducing or increasing the abundance of these structures are currently limited. We demonstrate that a long-acting form of IL-7 can be used to induce immune aggregates that include TLSs and increase the abundance of immune cells in the tumor microenvironment. This can be achieved without increasing the number of regulatory CD4 T cells or subsets of the myeloid population such as MDSCs. *Ltb*, a critical cytokine that regulates lymphoid neogenesis(31, 32) is enriched specifically in the TLSs induced by NT-I7. This finding is consistent with IL-7’s ability to enhance the survival of LTi cells and T cells that can produce LTβ(4, 11, 35). As such, NT-I7-mediated formation of TLSs in our model may be driven through LTi cells and the upregulation of *Ltb*.

The increase in the proportion of stem-like CD8 T cells after NT-I7 therapy is an intriguing finding. Stem-like CD8 T cells, also referred to as progenitor exhausted CD8 T cells, have a central memory-like profile and express the IL-7 receptor(36). IL-7 is an important cytokine for memory T cell formation and survival(10) that may be beneficial to stem-like CD8 T cells.

Preclinical studies demonstrate increased absolute numbers of stem-like CD8 T cells in MC38 tumors and increased proliferation in tumor draining lymph nodes after NT-I7 treatment(23). Our data suggest that NT-I7 therapy may support stem-like CD8 T cells in the tumor microenvironment. In addition to direct effects of NT-I7 on stem-like CD8 T cells, the long-acting form of IL-7 may also provide a protective niche for stem-like CD8 T cells by inducing the formation of TLSs in the tumor microenvironment. Stem-like CD8 T cells express CCR7 and CXCR5(36, 37), suggesting that they may preferentially home into lymphoid tissues. In support of this hypothesis, stem-like CD8 T cells are generally found in lymphoid tissues such as the spleen and lymph nodes in the chronic LCMV viral model that results in systemic viral infection(29). We have also demonstrated that stem-like CD8 T cells are preferentially located in tertiary lymphoid structures in human lung tumors(27). The results of our current study further reinforce the idea that stem-like CD8 T cells selectively home into lymphoid organs and tissues including TLSs within the tumor microenvironment.

Our results also suggests that there is a potential for synergy between NT-I7 therapy and immune checkpoint inhibitors. Enrichment of *Cd274* (PD-L1) *and Cd200*, the ligand for CD200R, indicate that the immune response within NT-I7-induced TLSs in the tumor microenvironment may be vulnerable to negative regulation of T cell function. This finding, coupled with the increased proportion and localization of stem-like CD8 T cells within TLSs, may provide an opportunity to improve the effectiveness of ICI. The clinical efficacy of PD-1 targeted therapy is critically dependent on stem-like CD8 T cells(36–41). Blockade of the PD-1 pathway results in a proliferative burst and subsequent differentiation of stem-like CD8 T cells into effector cells(36–38). Thus, combining PD-1-targeted therapy with NT-I7 therapy could enhance the anti-tumor CD8 T cell response and improve the clinical response to cancer immunotherapy.

In summary, long-acting NT-I7 remodels the tumor microenvironment to drive the development of TLSs while also enhancing the abundance of stem-like CD8 T cells, conventional CD4 T cells, B cells, and dendritic cells. This remodeling of the tumor immune microenvironment is associated with improved survival in our murine lung tumor model. Spatial transcriptomic analysis characterizes TLSs induced by NT-I7 and reveals potential combinatory strategies that could further enhance the efficacy of cancer immunotherapy.

## ACKNOWLEDGEMENTS

The project was supported in part by the American Lung Association Catalyst Award (CA-826809) to RCO. Additional support for this project was provided by the NUSeq Core at the Robert H. Lurie Cancer Center (P30CA060553), and the Light Microscopy Imaging, Flow Cytometry, PQHS Biostatistics, and Tissue Resources Shared Resources of the Case Comprehensive Cancer Center (P30CA043703). The study design, data collection, and analysis were done independently of funders.

## ETHICS STATEMENT

All animal studies were performed under a protocol approved by institutional animal care and use committee, and the methods confirm to the standards of good research practice. The study does not involve human subjects.

## DATA AVAILABILITY STATEMENT

The data generated in this study will be made available upon request from the corresponding author. Raw data from spatial transcriptomics analysis were generated at the NUSeq Core at the Robert H. Lurie Cancer Center, Northwestern University Feinberg School of Medicine and will be deposited into the Gene Expression Omnibus (GEO) archives. Data generated and presented in the supplementary figure are available from NeoImmuneTech, Inc. Restrictions may apply to the availability of these data, which were used under license for this study. Data are available from the authors upon reasonable request with the permission of NeoImmuneTech, Inc.

