## Supplementary Figure 1 for "Efineptakin alfa (NT-I7) improves overall survival and induces tertiary lymphoid structures in murine lung tumors"

**Supplementary Figure 1. Lymphoid structures in lymph node and intraperitoneal CT26 tumors**

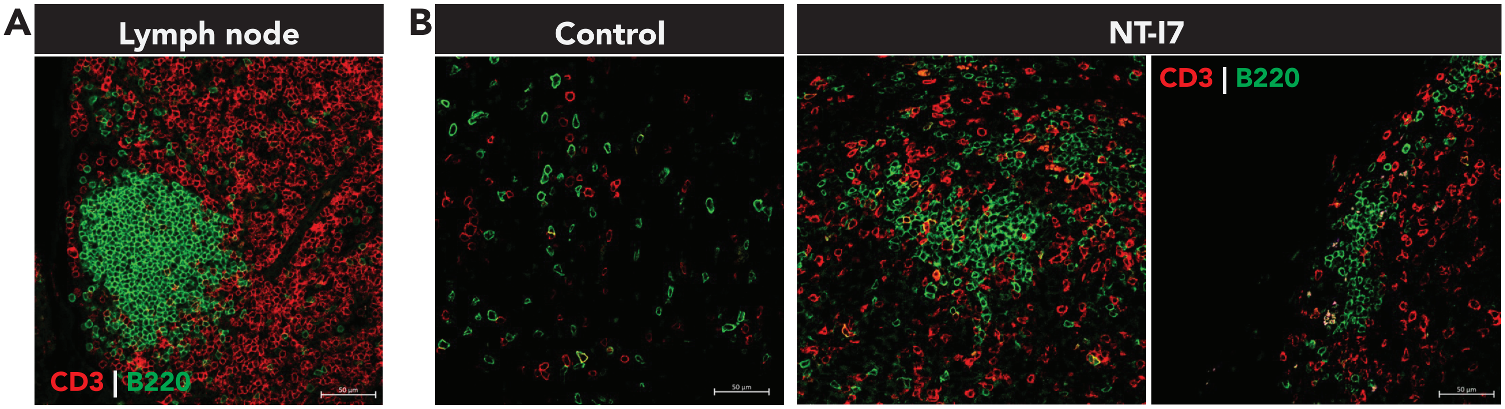

**Supplementary Figure 1. Lymphoid structures in lymph node and intraperitoneal CT26 tumors.** (A) Representative image of a conventional secondary lymphoid organ (peripheral lymph node) with classically organized T-cell/ B-cell compartments. Scale bar: 50 μm. (B) Left panel: representative image of i.p. tumor from control tumor. Right panel: representative images of individual i.p. tumors from the NT-I7-treated group. Scale bar: 50 μm. n = 1 for the control group and n = 3 for the NT-I7 treatment group.
